# VarQ: a tool for the structural analysis of Human Protein Variants

**DOI:** 10.1101/277491

**Authors:** Leandro Radusky, Carlos Modenutti, Javier Delgado, Juan P. Bustamante, Sebastian Vishnopolska, Christina Kiel, Luis Serrano, Marcelo Marti, Adrián Turjanski

## Abstract

Understanding the functional effect of Single Amino acid Substitutions (SAS), derived from the occurrence of single nucleotide variants (SNVs), and their relation to disease development is a major issue in clinical genomics. Even though there are several bioinformatic algorithms and servers that predict if a SAS can be pathogenic or not they give little or non-information on the actual effect on the protein function. Moreover, many of these algorithms are able to predict an effect that no necessarily translates directly into pathogenicity. VarQ Web Server is an online tool that given an UniProt id automatically analyzes known and user provided SAS for their effect on protein activity, folding, aggregation and protein interactions among others. VarQ assessment was performed over a set of previously manually curated variants, showing its ability to correctly predict the phenotypic outcome and its underlying cause. This resource is available online at http://varq.qb.fcen.uba.ar/.

Supporting Information & Tutorials may be found in the webpage of the tool.

## Introduction

The potential for genomics to contribute to clinical care has long been recognized, and the clinical use of information about a patient’s genome is rising. In this sense, precision medicine initiatives for disease treatment and prevention that take into account individual variability in genes are being implemented worldwide. Single nucleotide variants (SNVs) that manifest as protein variants are the most important form of variation in the genome, therefore a critical problem is to understand how single aminoacid substitutions (SAS) affect protein function and protein interaction networks (Vidal et al., 2011; Hu et al., 2016).

There are a several SAS effect predictors which perform a bioinformatic analysis and provide a pathogenicity score. Most of them are based on sequence information focusing on residue conservation and lacking a structural viewpoint which has proven to be critical to identify their effect (Kiel and Serrano, 2014). Moreover, even if they incorporate structural data (Bromberg and Rost, 2007; De Baets et al., 2012), usually they do not provide information that helps understand the molecular Moreover, even mechanisms underlying their prediction, thus preventing a personal assessment. This black box prediction is possibly rooted in the fact that proper (structural) analysis of possible effects of a given variant is time-consuming and requires expert handling of different tools, thus preventing its wide applicability in clinical genomics. As filtering algorithms allow researchers to identify a handful of variants that are likely pathogenic, manual curation and careful analysis are usually needed. Furthermore, even if a variant has been identified, mainly because it has been previously associated with disease, its effect on protein function could still be unknown.

Here we present VarQ, a Web Server that automatically analyzes several effects of SAS on protein function, particularly protein folding, activity, protein-protein interactions, and drug or cofactor binding. Analyzed variants are either automatically extracted from clinical databases and/or can be submitted directly by the user. VarQ automatically selects a critical set of representative structural configurations of the desired gene and diagnoses variants effect based on their impact. For this sake several properties are computed for each variant, including involvement in ligand binding, presence in the catalytic site (Porter et al., 2004), presence in protein pockets using the Fpocket software (Schmidtke et al., 2010), the free energy change using FoldX (Schymkowitz et al., 2005), the residue sidechain exposure to solvent, presence in a protein-protein interface as identified in the PDB’s structure and in the 3did database (Stein et al., 2009), the conservation of each residue in the Pfam family (Bateman et al., 2004), the switchability propensity using abSwitch software (Diaz et al., 2014) and the aggregability factor using Tango software (Fernandez-Escamilla et al., 2004). VarQ is intuitive, user-friendly, and provides clinicians, biochemists, geneticist and all professionals involved in personalized medicine with a straightforward tool to annotate and analyze protein variants. A detailed description of the programs and databases used is given in the Web Site.

## Methods

### Implementation of the tool as a bioinformatic pipeline

The VarQ tool is built around a collection of structural analysis tools tied together with the help of the workflow system Ruffus (Goodstadt, 2010). Having as unique input a list of UniProt (Magrane and UniProt Consortium, 2011) accession codes, the pipeline performs all the calculation steps described in Figure 1, parallelizing the acquisition of independent properties to improve its computational performance. Accession codes already computed are stored in a database to retrieve results without re-computing (this option works only for database stored substitutions and not for new user uploaded mutations).

**Figure 1:**
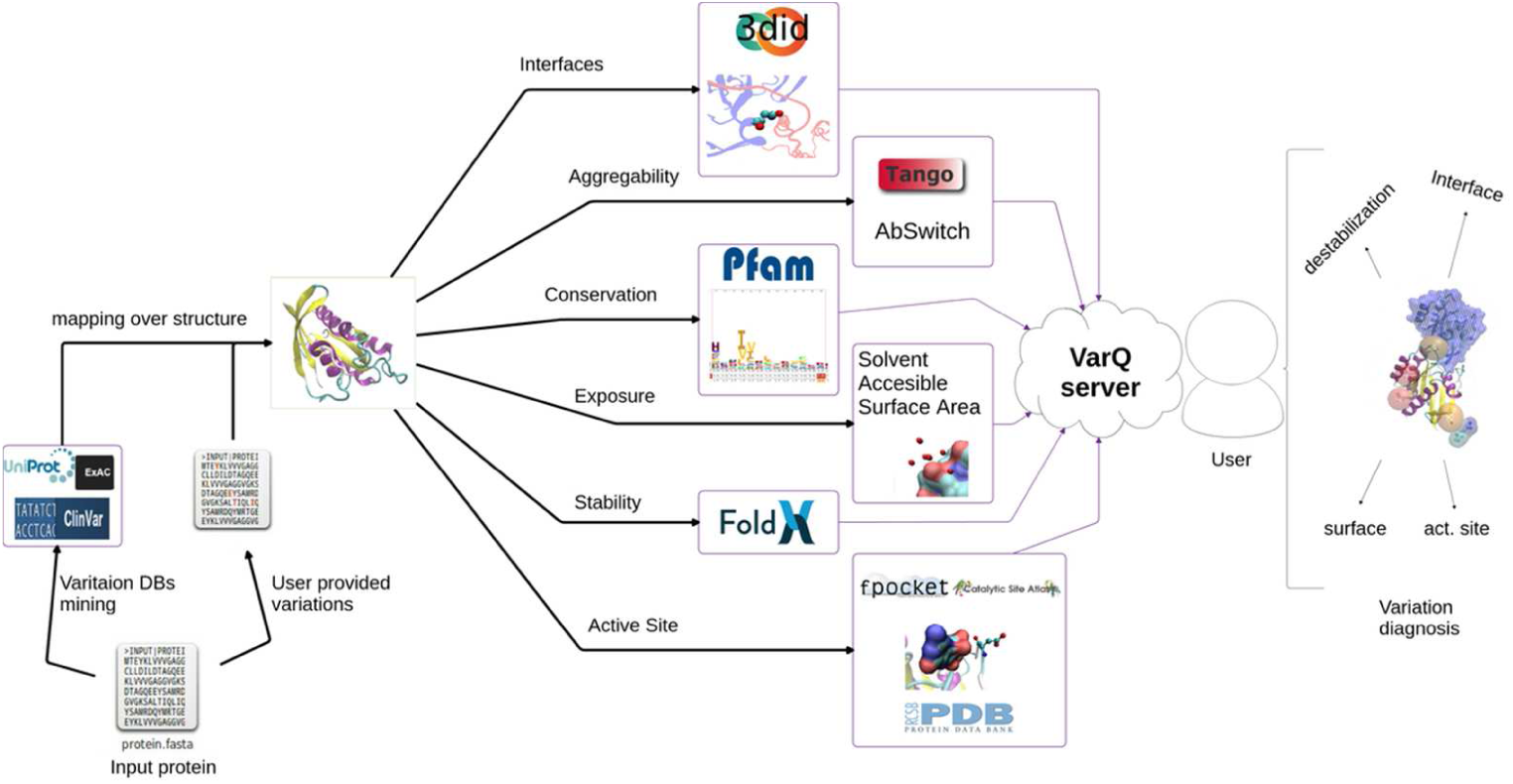
The VarQ general pipeline. Each known structural conformation for the input protein is analyzed independently to aid the user in the variant effect interpretation.

A Web Server developed with the Bottle Python library (bottlepy.org) allows both to visualize the results and download them for further analysis. Some of the features available are the mapping of the found variants on the protein sequence (Figure 2) and structure using JSmol (Hanson et al., 2013).

**Figure 2:**
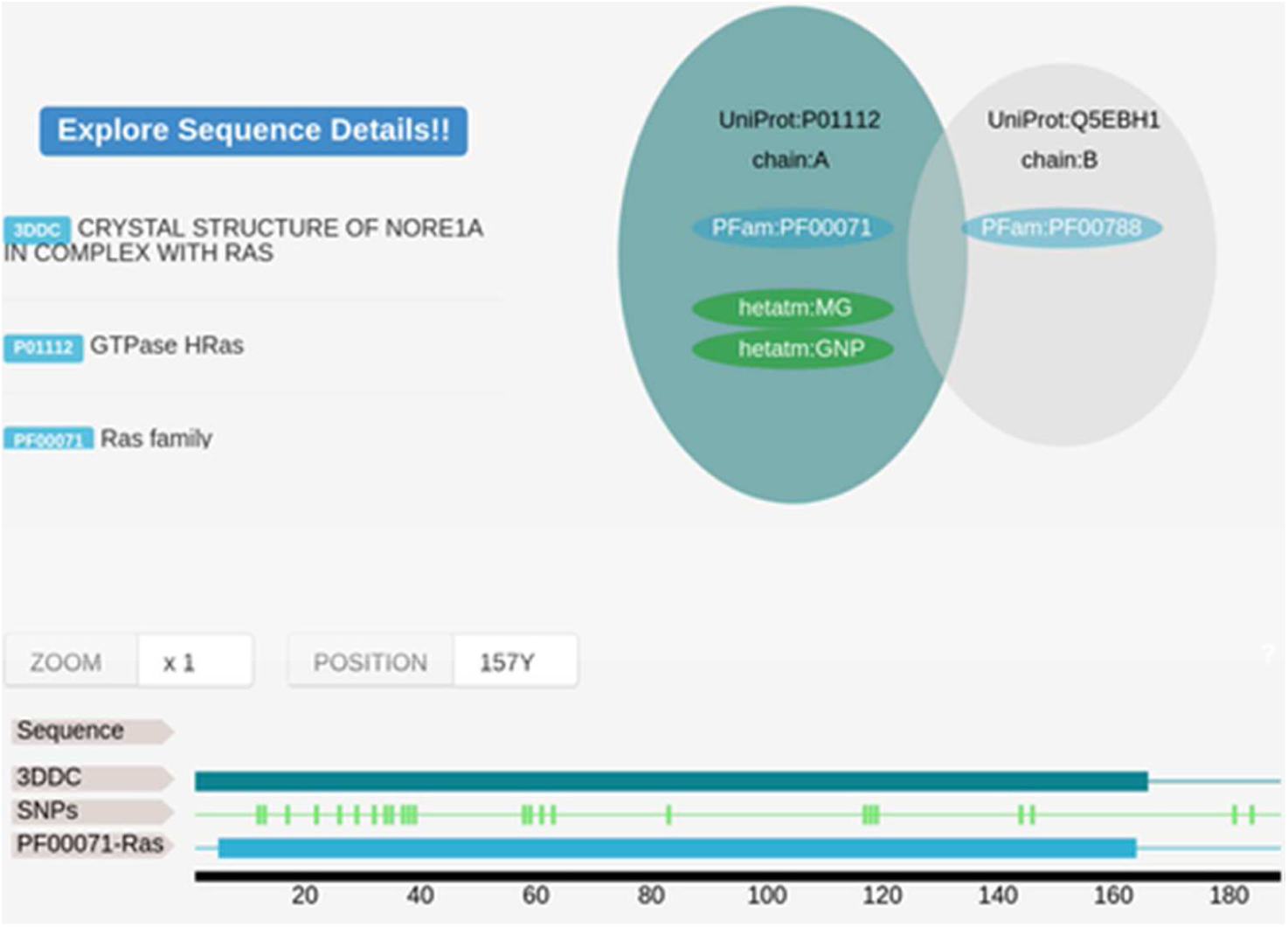
Sample VarQ output for the HRAS gene (PDB id 3DDC). On the left top, we show the structural features from crystallographic data. Target gene and possible interacting genes (when target gene was co-crystallized with other protein) are shown in green and grey respectively. The coverage of the crystals, the PFam families and the location of the variations are mapped over the UniProt original sequence.

### Structure Mining

When the user specifies a UniProt accession, the VarQ Pipeline searches within all the available structures that covers different segments of the sequence, preferring those with more coverage first and those with the best resolution in second place as a tiebreaker. Only those crystals covering at least 20 amino acids of the sequence of the target protein are considered. If there is more than one crystal structure covering the same part of the protein sequence, but coupling with different ligands, then both will be considered. If the same protein is crystallized in complex with another protein it will be considered separately and will appear with the information of the partner protein. The UniProt database is requested online to list all available crystal structures.

If the protein has not resolved structures in the PDB, an error message will be displayed in the web page together with 2 links to aid the user to search for possible solutions to this problem: one is the search of the UniProt accession in the RCSB web page and the second is the search of all crystallized UniProt accession codes that have the same gene name.

### Variation Mining

In this work we mine the variants of each Uniprot. Actually, there are several databases populated with variants coming from different sources: clinical trials (Landrum et al., 2016), sequencing information (Forbes et al., 2011), user submission (Day, 2010), etc. In this work, we are considering as a source of variants the following databases: i) UniProt annotated variants coming from dbSNP and BioMuta databases. ii) UniProt curated variants for human genes, UniProt database provides a database called humsavar (Famiglietti et al., 2014) that contains qualitative information classifying for all listed variants if they are disease-related or not and which is the disease involved in each mutation. iii) ClinVar variants provide additional mutations coming from clinical studies. When a protein target is introduced to our pipeline, all this information is read and the variations are kept for posterior analysis.

### Binding and catalytic residues

Each structure of the Protein Data Bank provides as an annotation for each crystallized ligand (compound, ion, cofactor, etc.) A dataset of solvent molecules were built to consider as binding residues only those interacting with non-solvents as binding residues. The information is parsed directly by the pipeline and the corresponding residues are labelled as involved in binding.

Also, we used the FPocket software for each protein structure considered in the analysis to calculate the protein pockets. We only considered the pockets with Druggability Score greater than 0.5, and/or if a ligand is present in the pocket. All the residues belonging to the pocket (even those not in contact with the ligand) are considered as binding residues.

The Catalytic Site Atlas (CSA) is a database documenting the active sites and catalytic residues of enzymes. CSA contains 2 types of entries: original manual-annotated entries, derived from the primary literature and homology-determined entries, based on sequence comparison methods to one of the original entries. All the residues belonging to the CSA database are labeled as catalytic residues in the VarQ pipeline.

### Changes in protein stability

All the mutations that were mapped to any protein structure were analyzed with the FoldX software. The FoldX software predicts the free energy change of a given mutation on the protein stability or on the stability of a protein-protein complex. Mutagenesis was performed using the BuildModel option of FoldX. The stabilities were calculated using the Stability command of FoldX, and ΔΔG values are computed by subtracting the energy of the WT from that of the mutant. The FoldX energy function includes terms that have been found to be important for protein stability. The equation describing the calculation of free energy of unfolding (ΔG) of a target protein is described in detail in the FoldX webserver. Briefly, it consists of an empirical force field that estimates the difference in the Van der Waals contributions of the protein with respect to the solvent, the differences in solvation energy for apolar and polar groups, the free energy difference between hydrogen bond formation, the difference in electrostatic interactions and the change in entropy due to fixing of the backbone and the sidechain in the folded state. When protein-protein stability is estimated, the empirical force field also includes a term related to the change in translational and rotational entropy and a term that takes into account the role of electrostatic interactions on the association constant. Those mutations that have more than 2 kcal/Mol are labelled as “High ddG variation”.

### Residue exposure

For each residue in each structure, the Solvent Accessible Surface Area (SASA) was computed The sidechain exposure percentages of each residue are informed, and those with more than 50% of its surface exposed are are labeled as exposed, or on the contrary, as buried. For this computation, the PyMol software (DELANO and L, 2002) is used as a command line tool calling to the get_area feature. The glycine residues are never labeled as buried or exposed since always present a 0% of exposure because of its absence of a sidechain. In the structural window, the value is displayed and a horizontal bar chart is shown based on the total side chain surface.

### Protein-Protein Interfaces

In order to detect those mutations that can affect protein-protein interaction, we decided to label the amino acids present in protein-protein interaction surfaces. To label a residue belonging to a protein-protein interface we evaluated the structures that have more than one chain. When an atom of a residue of one chain is at a distance of less than 5Å from an atom of the other chain the residue is labeled as interface residue. We also added an extra criteria; if the residue in the 3did database has been labeled as interacting we labeled the residue as 3did but we did not add the interface label. Despite the fact that the 3did database is not accurate enough for our automatic analysis pipeline, we decided to add this information since we considered it valuable to evaluate possible effects as it expands the database of residues which are involved in protein-protein interfaces, but needs manual inspection.

### Other properties

The BFactor of a residue usually relates to its mobility and we inform this value with respect to the median of the B-factor of all the residues in the protein. We also calculate the Switchability, which informs the tendency of residues to switch from alpha-helix to a beta sheet type of secondary structure; the Aggregability, which is the tendency of a residue to generate aggregation when is mutated and is computed using the Tango Software; the conservation, since conserved positions are expected to be relevant for protein function, calculated as the value in bits of the corresponding letter of the original amino acid in the Pfam family only if it is assigned for the analyzed position. This value is computed by an in-house developed script.

## Results

The web server is developed to run proteins based on its UniProt ID. To test our developed pipeline we first run a manually curated set of variants derived from our previous work (Kiel and Serrano, 2014), comprising mutations in Ras/MAPK pathway components which can cause either cancer or developmental a group of disorders called RASopathies (Tidyman and Rauen, 2009). This previous case by case analysis required a considerable amount of manual curation and now constitutes an excellent benchmark for our web server capabilities. The results obtained with the newly developed pipeline (Figure 1) as implemented in the VarQ web server where all the calculations and decisions run fully automatic are presented in Table 1.

**Table 1:** Comparison of mutation-mining of previous hand-curated work against the results given of VarQ Pipeline for benchmark set of RASopathies related proteins.

|  | Kiel & Serrano 2014 | VarQ |
| --- | --- | --- |
| Total variants | 956 | 1109 |
| Mapped onto structure | 624 | 566 |
| Predicted to be neutral | 197* | 152 |
| Predicted to be affect protein function | 427* | 414 |

* In Kiel & Serrano 2014 variants were classified as neutral or disease causing based on FoldX energy score.

Figure 2 shows an example of the VarQ output for the HRAS gene (PDB id 3ddc) depicting the crystallographic unit and the retrieved variations mapped in the protein sequence. PFam family information and mapping within protein sequence is also shown.

VarQ automatically retrieved 566 variants, out of the 624 mutations identified manually and structurally mapped by Kiel and Serrano (Table 1). From these, we were able to automatically identify 414 (≈ 97% efficiency) of the 427 pathogenic variants.

Figure 3 shows the histogram of the ΔΔG energy, estimated using the FoldX software, for neutral and pathogenic SAS. As clearly evidenced, a value higher than 2 kcal/mol is a good indication that the SAS and thus, the underlying mutation is pathogenic, However, for lower ΔΔG energy values, other properties need to be considered to be able to define potential pathogenicity and also to understand the underlying molecular mechanism leading to disease.

**Figure 3:**
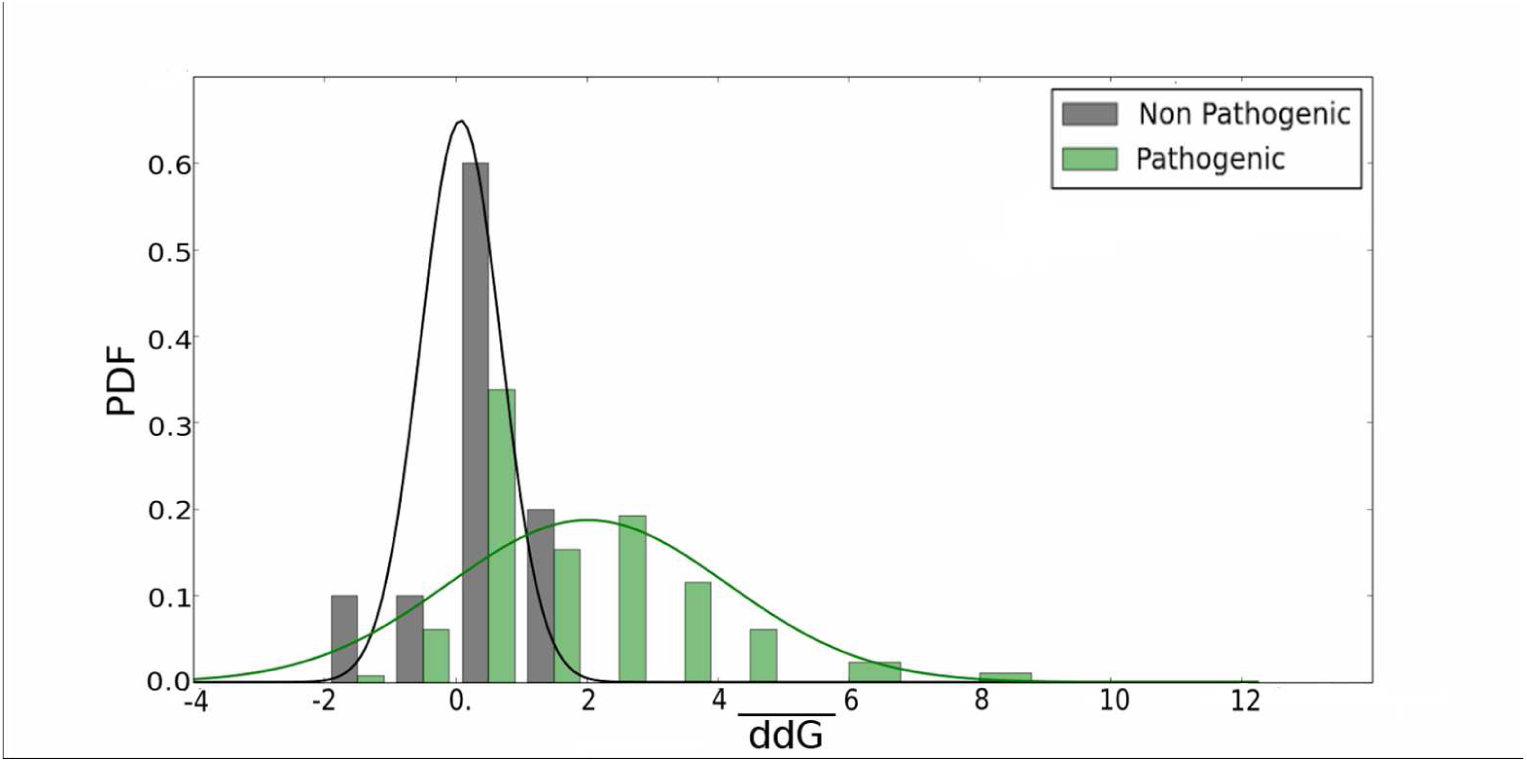
Energy variation of the pathogenic-labeled mutations versus the non-pathogenic-labeled mutations computed by the FoldX software for the set of mined mutations over the RASopathies related proteins.

VarQ pipeline, further classifies all the amino acids of each protein automatically, in key categories related to their function (being part of the active site, of protein-protein interaction surfaces, etc. see methods for details) and presents that information to help decision making (Table 3). For example, we were able to identified 94% of the manual curated protein-protein interaction variants; also all variants with high switchability and aggregability, which were correctly classified as folding disruptors. The active site and binding residues are those either labeled as ligand-binding residues in the PDB file, those belonging to the same pocket of these residues as determined by the Fpocket program, and those belonging to the Catalytic Site Atlas database. For example, there are two HRAS structures, binding GNP (i.e. PDB id 3ddC) and GTP (i.e. PDB id 4k81), and reported mutations in the binding pocket have been proposed to modify signalling cascades which are involved in different diseases (Prior, 2012). VarQ properly labeled these residues as active site residues in each structure, thus allowing the user to diagnose SAS effects in those signalling pathways.

Folding residues are those not inter-domain and not Active Site having an accessible surface area percentage lower than 50%. In the particular case of the RASopathies proteins, a high proportion of the residues are marked as interface-involved because these proteins participate in a very complex network with a large set of protein-protein interactions. Localization mutations (a total of 3, see Table 2) are not included in our analysis.

**Table 2:** Classification of the previous work (FoldX destabilizing/disease-causing mechanism know category) compared with the automatic classification applied to VarQ results. Our method do not label any mutation as “localisation” so we kept the label used by Kiel & Serrano. Number in parenthesis are the percentages of the total residues with that label.

| Effect in | Kiel & Serrano 2014 | VarQ |
| --- | --- | --- |
| Active site + Binding | 200 (47%) | 203 (49%) |
| Protein-protein interaction | 79 (18%) | 74 (18%) |
| Folding | 145 (34%) | 137 (33%) |
| Localisation | 3 (1%) | - |
| Total | 427 | 414 |

**Table 3:** Decision tree to propose a possible effect of mutation in an analized protein.

| $\Delta\Delta G$ | Active site | Interface | Buried | High aggregability | High switchability | Diagnostic | Website Outcome Label |
| --- | --- | --- | --- | --- | --- | --- | --- |
| ↑ | ✓ |  |  |  |  | Protein activity altered | Disrupt protein function |
|  | ☐ | ✓ |  |  |  | Protein-protein interaction affected | Disrupt protein interface |
|  | ☐ | ☐ | ✓ |  |  | Destabilization of domain preventing folding | Disrupt protein folding |
|  | ☐ | ☐ | ☐ | ✓ |  | Destabilization of domain preventing folding | Disrupt protein structure |
| ↓ | ✓ |  |  |  |  | Potential protein activity altered | Potential disruption protein function |
|  | ☐ | ✓ |  |  |  | Potential protein-protein interaction affected | Potential disruption protein interface |
|  | ☐ | ☐ | ✓ |  |  | Potential destabilization of domain preventing folding | Potential disruption protein folding |
|  | ☐ | ☐ | ☐ | ✓ |  | Potential destabilization of domain preventing folding | Potential disruption protein structure |
| ↓ | ☐ | ☐ | ☐ | ☐ | ☐ | No effect | Likely non pathogenic mutation |

In summary our pipeline was able to identify automatically most of the previously identified variants (Kiel and Serrano, 2014), which were correctly classified according to their function and localization in the protein. This statistics are summarized in Table 2. Based on that information and our bioinformatic analysis, well-defined and relevant effects were proposed for each variant. Due to the inherent complexity of the problem, we believe that an automatic pathogenicity prediction cannot be offered but the automatic classification and information provided by VarQ can be used to help decide the possible effect of a mutation and therefore help scientists in their interpretations and pathogenicity evaluation.

Within the web page, users can find detailed tutorials explaining how to load new jobs in the server, a detailed explanation of the output obtained with each job and how to interpret results in order to diagnose the SAS effects in the protein structure.

## Discussion

In this work we introduce VarQ, a novel online tool that offers an user friendly way to evaluate the effect of protein variants that arise from human genomics projects. We have shown that VarQ is able to correctly reproduce previous analysis of RASopathies related mutations avoiding extensive and time consuming manual curation. Also, we assigned 153 new mutations that represent novel cases that cannot be compared with the previous study.

Users can use VarQ to either analyze mutations that are already available in clinical databases or to analyze novel unreported variants. To assist variation diagnosis each analyzed mutation is labeled according to several computed properties. For different conformations of the same protein (i.e. active and inactive determined structures) the same mutation could lead to different labeling, leaving to the user the final assessment of the effect caused by the variation. VarQ displays its full potential on human proteins with known structure(s) but can also be used with any protein. Usually clinical researchers, biochemists and geneticists do not have the bioinformatic resources to massively analyze variants so they use web servers or easy to use softwares to classify the variants. On the other hand, bioinformatic softwares usually are aimed at predicting only pathogenicity but do not give information for the users to be able to gain insight and finally decide the possible/potential effect of the mutation. This information is of paramount importance when pathogenic prediction is challenging. In this sense, for example a mutation that disrupts a protein-protein interaction may be pathogenic or could be benign depending on the protein function and the biology of the interaction. To the best of our knowledge this is the first application that provides this information in an automatic, simple and intuitive way.

